# Cognitive load dissonance and personality factors: an empirical analysis in organizational settings

**DOI:** 10.1101/2023.12.05.570254

**Authors:** Di Gruttola Francesco, Mastrogiorgio Antonio, Orfei M. Donata, D’Arcangelo Sonia, Lattanzi Nicola, Ricciardi Emiliano, Malizia P. Andrea

## Abstract

Previous research that aimed at characterizing the importance of workers’ personological traits in coping with stress in organizational settings is often biased by the potential inconsistency (i.e., dissonance) between subjective perception and objective experience of workload. This study explored the relationship between the subjective, self-reported and objective physiological measures of cognitive load, and the possible confounding role of personality traits on a representative population (call center operators) in ecological settings (daily working routines consisting in inbound and outbound calls). With this aim, the personality traits of 30 call center operators were preliminarily characterized using the Ten Item Personality Inventory. Then, objective heart rate variability and electrodermal activity were measured during their inbound and outbound calls. Finally, a subjective self-evaluation of the experienced cognitive load was acquired. No significant correlations were found between subjective and objective measures of cognitive load except when controlling for personality traits. In particular, a negative correlation was found between the subjective perception of cognitive load and the psychophysiological indices of the parasympathetic tone. Specifically, the personality factors of Openness to Experience, Agreeableness and Emotional Stability have a significant influence on the subjective perception of cognitive load, without predicting the objective psychophysiological expression. Our study emphasizes the importance of studying the dissonance between subjectively-perceived feelings and their objective physiological instantiation, along with the influence of personality factors, in organizational settings.

## 1. Introduction

The psychophysiological characterization of *cognitive load* (hereafter, CL) is a classical topic of organizational research, mainly analyzed with reference to workload and stress in workplaces (e.g., Hancock & Warm, 1989; Hockey, 1997; Lundberg & Frankenhaeuser, 1999). Though the involved physiological mechanisms are relatively known, the phenomenon is not trivial as it is affected by a number of strictly psychological factors. A number of studies investigated the role of personological traits in influencing the psychophysiological response to CL related to working activities. Relatively to a personality model widely studied for its association with CL, the Big Five personality traits — Neuroticism, Openness, Agreeableness, Extraversion, Consciousness (Costa & McCrae, 1992) — the evidence is markedly mixed (e.g. Pollack et al., 2020; Xin et al., 2017).

Understanding the reasons for such mixed results is not an easy task. Overall, previous research, focusing on the substantive problem of stress in the workplace, discharges a more fundamental matter: the potential inconsistency between *subjective*, self-reported CL and its *objective* physiological dimension, as indexed by measures like heart rate variability, pupil size and electrodermal activity. Indeed, the implicit assumption in literature is that a subjective perception of CL — what an individual *subjectively* perceives — is a reliable and accurate proxy of the objective physiological response — what an individual *objectively* experiences. But this implicit assumption is precisely the one, at stake, we aim to investigate. Indeed, an eventual *dissonance* between subjective perceived CL and its objective physiological instantiation could jeopardize the validity of personological traits in influencing the CL. In addition, this dissonance could matter in the analysis of other typical constructs, such as health, job performance and burnout, potentially related to CL. Experiencing stress while reporting a calm situation is just one of the marks of dissonance, representing a risk factor for workers’ well-being.

In this study, we explored the relationship between the subjective and objective measures of CL, and assessed to whether and to what extent such a relationship is confounded^1^ by personological traits (see Fig. 1A). In order to analyze such a relationship, we deliberately opted for organizational settings. We conducted our analysis I) on a *representative population*, consisting of call center operators, and II) in *ecological settings,* consisting of daily job routines represented by inbound and outbound calls. Our study emphasizes the relevance of investigating the consistency between subjective perceived feelings and objective physiological responses, along with the confounding role of personological factors, on a population of real workers in ecological settings.

**Figure 1:**
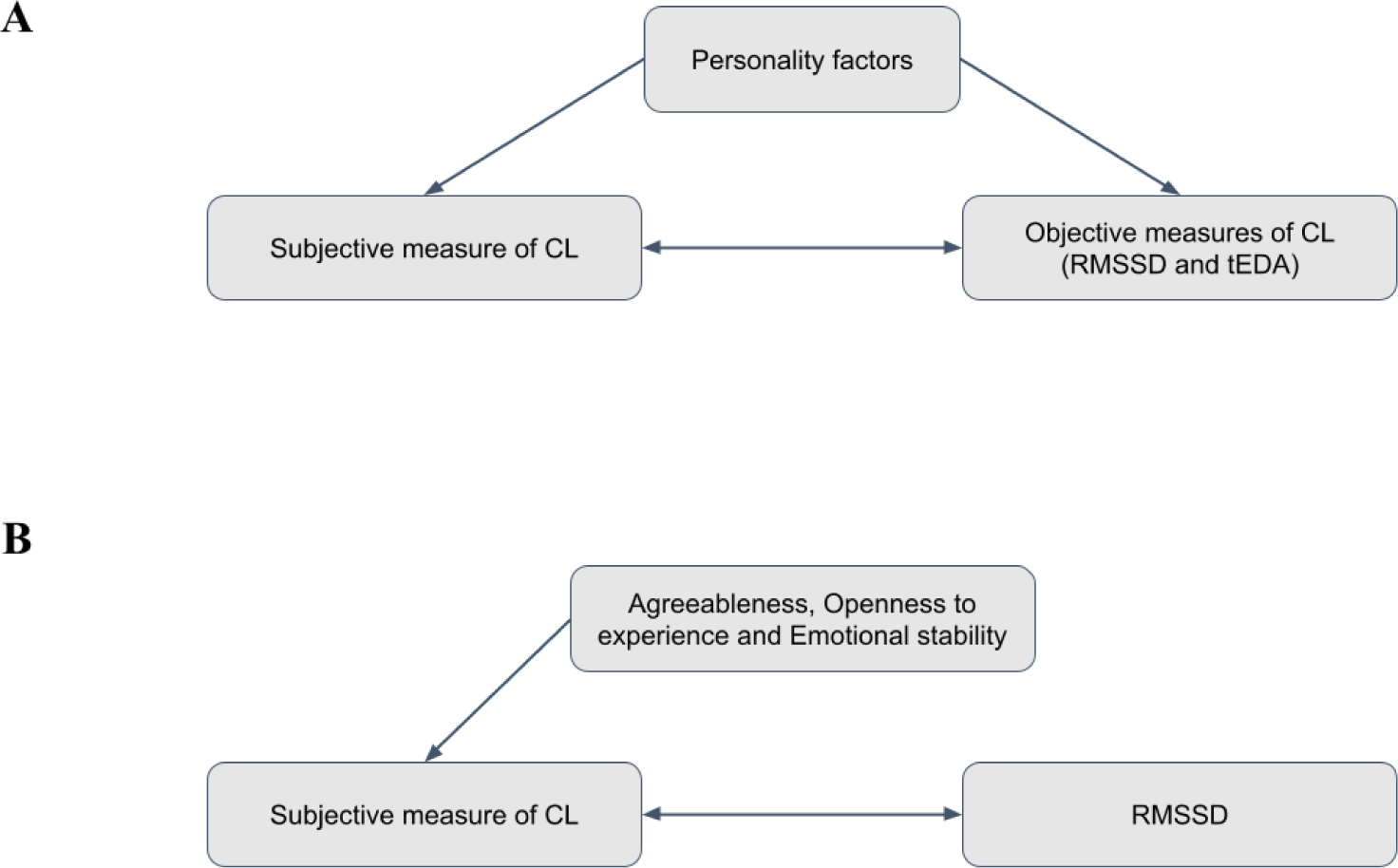
Our Hypothesis 2 is represented. According to the literature, personality is independently associated both to subjective and objective measures of cognitive load. Thus, we hypothesized a confounding role of personality factors in the relation between subjective and objective measures of cognitive load, possibly influencing the direction and size of the correlation (A). We actually found a causal association of Agreeableness, Openness to experience and Emotional stability on the subjective measure of cognitive load. Thus, we could theorize a possible confounding role of these three personality dimensions on the relation that we found between participants’ self-assessment of cognitive load and the vagal tone index (RMSSD). Specifically, Agreeableness, Openness to experience and Emotional stability could indirectly influence this correlation by affecting workers’ subjective perception of cognitive load (B).

### 1.1 Theoretical background

The classical CL theory, referring to Baddeley’s model of working memory (Baddeley & Hitch, 1974), assumes that the human mind has limited cognitive capabilities in storing information and in paying attention to multiple stimuli (Klepsch, Schmitz & Seufert, 2017; Skulmowski & Rey, 2017). Thus, the CL of a given task may depend on its complexity (intrinsic CL), on the difficulty in finding or effectively understanding the information required to perform it (extraneous CL), or on the possibility to use strategies to facilitate the task or empower our cognitive capacity, such as taking notes (Germane CL - Klepsch, Schmitz & Seufert, 2017; Skulmowski & Rey, 2017). A *subjective* measure of CL is typically detected through self-reports or subjective questionnaires aimed at investigating participants’ mental effort in performing different kinds of tasks (Klepsch, Schmitz & Seufert, 2017; Skulmowski & Rey, 2017). On the other hand, the *objective* physiological responses related to CL are assessed through multiple measures that may include heart rate variability, pupil size and electrodermal activity, able to detect participants’ sympathetic and parasympathetic autonomic nervous system (hereafter ANS^2^) responses during the task (see Forte & Casagrande, 2019; Massaro & Pecchia, 2019; Wang et al., 2018; Craft & Schwartz, 1995; Sugenoya et al., 1990).

Personality could interfere with CL by influencing both the subjective perception and the objective physiological responses. Multiple studies found a general influence of personality on psychophysiological response to workload in stressing conditions. With reference to the Big Five personality traits — Neuroticism, Openness, Agreeableness, Extraversion and Conscientiousness (Costa & McCrae, 1992) — Xin et al., (2017) found that Neuroticism, Extraversion and Openness predicted the heart rate stress reactivity, the cortisol response and the affective evaluation of participants who had been experimentally induced through a stressful situation. Similarly, Pollak et al., (2020) underlined how Emotional Stability, Openness to Experience and Conscientiousness, predicted stress appraisal of students enrolled in an industrial robot programming course. However, when focusing on the influence of each personality trait, the evidence is mixed as different studies reported contradictory results, in both size and direction of the effects: in some studies, a specific trait magnifies the psychophysiological response to workload, in others the same one exerts a negative or null effect (Bibbey et al., 2013; Garcia-Banda et al., 2011; Oswald et al., 2006; Xin et al., 2017). In fact, with reference to Openness to experience, previous studies emphasize greater stress resilience among open individuals (Williams et al., 2009; Lü, Wang & Hughes; 2015), while others show opposite outcomes (Bibbey et al., 2013; Oswald et al., 2006). Conscientiousness has been reported to be associated with higher physiological activation in stressed condition as “is one of the personality dimensions more strongly related to health status perhaps partly because such individuals display good self-control, responsibility, and prosocial attitude towards others” (Garcia-Banda et al., 2011, p.1345), while no association has been reported other studies (Oswald et al., 2006). Regarding Neuroticism, a higher level of this trait has been associated both with high level of perceived work-related stress (Gramstad, Gjestad & Haver, 2013) and, in certain conditions, even with lower stress reactivity according to the emotional or cognitive nature of the proposed task and to participants’ gender (Jonassaint et al., 2009; Oswald et al., 2006), although several studies did not report associations (Kirschbaum et al., 1995; Schommer, et al., 1999; Vassend & Knardahl, 2005). For Extraversion, there is no strict evidence of a moderation role (Schommer et al., 1999; Vassend & Knardahl, 2005), although some studies report that higher extraversion was linked to lower reactivity (Jonassaint et al., 1999). Finally, with reference to Agreeableness, null evidence has been found (Oswald et al., 2006; Williams et al., 2009).

The reasons for such mixed results are unclear. As claimed by Bibbey et al. (2013, p. 29), “Given the paucity and inconsistency of previous research, we had no clear expectations regarding the size and the direction of any association between stress reactivity and the other personality traits that make up the Big Five” (see also Kirschbaum, Pirke & Hellhammer, 1993).

### 1.2 Open issues in organizational settings

Previous studies have often been conducted in poor organizational settings, that is, they have often been conducted on *non-representative populations*, such as graduate and postgraduate students (e.g., Birkett, 2011; Kirschbaum, Pirke & Hellhammer, 1993; Verschoor & Markus, 2011; Von Dawans, Kirschbaum & Heinrichs, 2011) or in *non-ecological conditions*, such as learning environments (Mutlu-Bayraktar, Ozel, Altindis & Yilmaz, 2020), often through laboratory-induced extreme conditions, such as acute stress. In this theoretical framework, limited evidence about the relation between subjective and objective measures of CL is available (Makransky, Terkildsen & Mayer, 2019; Minkley, Xu & Krell, 2021; Mutlu-Bayraktar, et al., 2020).

Non-representative populations and non-ecological conditions represent recurrent features of previous studies. Such features jeopardize the possibility of evaluating the consonance — and the eventual dissonance — between subjectively perceived and objective experienced CL, and to investigate the confounding role of personality.

#### Population representativeness

Existing studies (mentioned in the previous 1.1 section) have often been conducted on non-representative populations, such as graduate and postgraduate students (e.g., Verschoor & Markus, 2011). This condition represents a fundamental limitation for external validity as nothing grants that existing evidence can be generalized to organizational populations. External validity is not an ancillary matter: CL is a theoretical construct that presents substantial specificities when analyzed in organizational settings. In particular, workers’ well-being is a topic of theoretical and applicative relevance for Industrial and Organizational Psychology. This is even more considerable if we refer to professions whose conditions or type of work make them more prone to develop physical and psychological issues.

In this regard, one of the most vulnerable categories are the call center operators (Conway et al., 2013; Jeyapal, Bhasin, Kannan & Bhatia, 2015; Oh, Park, & Boo, 2017) because of the low salary, precarious job status, stressful work setting and - consequently - the high risk of physical and psychological problems, such as anxiety, depression, stress and burnout (Conway et al., 2013; Costa & Costa, 2017; Jeyapal et al., 2015; Oh et al., 2017; Rameshbabu, Reddy & Fleming, 2013). In this vein, Toker and Güler (2021) reported that at least 45.8% of call center operators have vulnerable mental health. In addition, call center operators make extensive use of technological devices, especially computers and telephones, to interact with clients. Their daily activities are mainly divided into outbound and inbound calls (Lin et al., 2010). Outbound calls are made by the call center operators to sell products or reach new clients. On the contrary, they receive inbound calls from customers to give information or to solve problems encountered in the use of company services. Both types of activities are reported to be a meaningful source of stress, especially when dealing with difficult or demanding customers (Conway et al., 2013; Lin et al., 2010).

#### Ecological conditions

The relationship between subjective perception and objective psychophysiological measures of CL has not been investigated in strict organizational environments. Indeed, existing research often relies on laboratory tasks. For instance the investigation on the relation between objective and subjective measures in call center operators (Enoki, Maeda, Iwata & Murata, 2017) was not performed during their typical working day activities. An artificial reproduction of an organizational setting can limit participants’ agency and embodiment by involving different neuropsychological mechanisms compared to an ecological context (Shamay-Tsoory & Mendelsohn, 2019).

Furthermore, the typical use of laboratory tasks is not just problematic because of the lack of ecological conditions but it is potentially “biased” as it focuses on extreme conditions. For instance the Trier Social Stress Test has been largely used for this purpose (Birkett, 2011; Von Dawans, Kirschbaum & Heinrichs, 2011) during which stress is induced through speech and arithmetic tasks to be accomplished in presence of an evaluative audience. Such studies focus on acute stress relying on experimental designs in which high vs low conditions are compared. In such studies, standard conditions (not abnormal ones) are, paradoxically, under-investigated. The comparison of extreme conditions, such as high vs low CL, is based on the assumption of a linear relationship between CL and psychophysiological response. Actually, the existence of U-shaped or phasic relationships is often overlooked along with their explanations involving hormesis-like phenomena (Calabrese & Baldwin, 2003) in which the response in correspondence of a low dosage can be (non just of different magnitude but) of different type in correspondence to high dosage. The use of extreme experimental conditions, in existing research, represents a further limitation to generalizability and a focus on non-extreme, standard everyday working conditions represents, paradoxically, a novelty.

Everyday tasks of call center operators represent a prototypical ecological setting as such tasks are instantiated in standardized, non-extreme, stable working activities. Research reports that call center operators experience episodes of dissonance on a daily basis (Molino et al., 2016; Zito et al., 2018). Here dissonance refers to the discrepancy between the subjective perception and the objective physical/behavioral expression of a given state (cf., Festinger, 2022/1957). For instance, call-center agents need to maintain proper tone, language and express positive emotions despite frequently experiencing frustration with customers or possible problems arising from their work and personal life. This continuous lack of consonance between subjective perception and objective expression of their emotions could be a risk factor for their well-being, affecting job satisfaction and turnover intentions (Zito et al., 2018).

## 2. Materials and Methods

### 2.1 Hypotheses

Considering the theoretical and methodological gaps of the literature (discussed in the previous sections) our aim is to study CL dissonance and the possible confounding role of personality traits in the population of call center operators involved in daily working routines represented by inbound and outbound calls. Studying the CL dissonance in such natural settings — a real organizational population observed in the ecological conditions of daily working activities — could reveal different psychophysiological underpinning.

Moving from such premises, here, we expect to find evidence of CL dissonance in call center operators (*Hypothesis 1*). In fact, this phenomenon is widely reported in the studies that consider this special population (Zito et al., 2018). Thus, a lack of correlation between the subjective and objective measures of CL — as an evidence of CL dissonance — is expected. Furthermore, we hypothesize that as in the general population, personality factors can also be a confounder in this relationship highlighting a correlation between subjective and objective response of CL (*Hypothesis 2.1*) and predict the latter variables (*Hypothesis 2.2 - Figure 1A*). In fact, a relationship between personality traits and CL - both subjective and objective - emerges in the literature, with the CL operationalized as a response to work-related or experimentally induced stress, and personality traits seem to be related both to a subjective assessment of the stressful event and to its physiological response (Xin et al., 2017; Garcia-Banda et al., 2011; Oswald et al., 2006). However, we do not advance precise hypotheses on the direction and magnitude both for its confounding effect and its predictive role on the subjective and objective measures of CL, because the evidence in literature is mixed (Bibbey et al., 2013) and to date this topic has never been investigated in an ecological setting, as proposed in our study.

### 2.2 Participants

A sample of 30 Italian bank workers (17 females, 13 males) with a mean age of 35 years (SD ± 11) whose occupation was to give technical and commercial support to their customers remotely through the phone joined the experiment. The recruitment period started the 8th of April 2019 and ended the 3rd of May 2019. Participants freely and voluntarily took part in the research through an internal call from their company by signing a consent form explaining the purpose of the research and the methods of processing personal data. Before the starting of the experimental phase, we personally explained again the aims of the study. Then, participants were assigned a progressive number from 1 to 30 so that we could not trace their identity. No sensitive data (e.g. participants’ personal details or contact details) were collected. The study was conducted in accordance with the ethical principles of the Declaration of Helsinki (World Medical Association, 2013) and under a protocol approved by the Area Vasta Nord Ovest Ethics Committee (protocol n. 24579/2018).

### 2.3 Procedure

The research was organized in two sessions. In the first, participants filled out a personality questionnaire via an online Google Form. The second part took place during working mornings and afternoons within participants’ ordinary workplace environment in three Italian customer care facilities of the bank (Torino, Milano and Bologna) where participants performed both inbound and outbound calls. In each location, the investigators prepared and controlled the experiment in a quiet and bright room away from workstations without interfering with workers’ jobs. Participants were previously instructed about the place and time of the experiment and, for each experimental session, two participants were tested in parallel. Firstly, they entered one by one in the investigators’ room, received the research instructions, and were fitted with a wristband that measured the heart rate variability, the body temperature, the motion-based activity and the electrodermal activity. The wristband was chosen as physiological device because of its lightness, wearability and low invasiveness in participants’ working activity. The device was placed on the wrist of the left hand, which was less involved in the work-related body movements of the participants compared to the right hand that was used to control the mouse of their PC. Then, participants’ resting psychophysiological activity was measured inside the room. This rest period served as baseline of participants’ starting physiological activity before recording it during working hours, thus making the measurements comparable across sessions and participants. Participants sat comfortably in a chair and were instructed to remain for two minutes in a static position, with their eyes open and without speaking. Then, they reached their workstation, and the psychophysiological correlates of their typical working routine were measured for two hours. The participants’ workstation was well lit and consisted mainly of a table at which they were seated and a PC with a mouse and a keyboard. Participants wore a microphone with headphones to talk to customers. During this period in case of technical problems or any breaks, the experimenters remained available inside the room. At the end of the two hours, each participant returned to the experimenters’ room and the wristband was removed. This device had a button that was used to mark the start and end of each evaluation phase (rest and experimental). Once the wristband was removed, a final question concerning the subjective perception of the CL experienced during the two hours of evaluation was made to each participant. Each participant was assessed for two days in a row, thus assessing 4 hours of daily working routine in total. For each day we followed the same protocol and scheduled the assessment of each participant during the same part of the day (morning or afternoon) in order to make the within-subject assessments comparable over the two days. The morning and afternoon assessment times were variable covering the initial, central and final time slot, thus having a complete coverage of all phases of the day. This modality fitted with the working need and the shifts of the participants, who had part-time or full-time contracts, therefore working mainly in a part (morning or afternoon) or the whole day. Considering the whole sample, the assessments were balanced with half of participants evaluated in the mornings and half in the afternoons.

### 2.4 Measures

#### 2.4.1 Personality traits

The Italian version of the Ten Item Personality Inventory (TIPI – see Chiorri et al., 2015a) was used to measure participants’ personality traits. This questionnaire is a short version of the Big Five Personality Inventory (BFI) with ten items in total. It was selected due to its quick administration time and for its good psychometric properties, showing an adequate factor structure, test-retest reliability, self-observer agreement and convergent and discriminant validity with the BFI (Chiorri et al., 2015a). Participants were asked to evaluate whether each presented trait may or may not have been applied to them by rating the level of agreement on a scale from 1 (disagree strongly) to 7 (agree strongly). The scale measures the presence of five personality factors (two items per factor) with higher scores corresponding to a higher presence of the factor itself:

1. Extraversion: from being reserved and quiet to extraverted and enthusiastic;
2. Agreeableness: from being critical and quarrelsome to sympathetic and warm;
3. Conscientiousness: from being disorganized and careless to dependable and self-disciplined;
4. Emotional Stability: from being anxious and easily upset to calm and emotionally stable;
5. Openness to Experiences: from being conventional and uncreative to open to new experiences and complex.

Firstly, we computed the score of the reversed items (one for each personality factor). Then, we calculated the score of each personality trait taking the average between their two corresponding items (Chiorri et al., 2015a; Gosling, Rentfrow & Swann, 2003).

#### 2.4.2 Heart Rate Variability

The Heart Rate Variability was measured with the Empatica E4 wristband (Empatica S.r.l.). This device was equipped with a Photoplethysmography (PPG) sensor that measured the Blood Volume Pulse (BVP) with a sampling rate of 64 Hz. For each day of evaluation and for each participant, considering the duration interval between adjacent BVP peaks, the inter-beat-interval (IBI) in milliseconds of the entire time window of registration and the Heart Rate (HR) were computed. These data were exported in CSV format with the E4 manager software (Version 2.0.3.5119) and analyzed through the Kubios HRV Standard software (Version 3.4.3). Each HRV signal was divided into two epochs of 2 and 120 minutes, that is the rest and the experimental condition, respectively. Each epoch was independently inspected for artifacts (i.e., missing or extra beats) both visually and with a threshold-based artifact correction algorithm. The latter algorithm, within each epoch, compared each IBI against the local average interval filtered by the median of the entire time window. For each epoch, an HRV expert in artifact detection marked an IBI as artifact both when the Beats Per Minutes (BPM) corresponding to the IBIs were outside the typical physiological range (below and above 3 SD from the mean) and the distance between the IBI and the local average was higher than 0.45 seconds (Laborde, Mosley & Thayer, 2017). Then, the artifactual IBIs were removed and replaced, together with the missing IBIs, via a cubic spline interpolation algorithm. Then, time domain and non-linear measurements were computed on each epoch. The root mean square of successive differences between normal heartbeats (RMSSD), computed as the average of the IBIs square root, was obtained as an index of the vagal tone (Laborde et al., 2017; Shaffer & Ginsberg, 2017). The higher the RMSSD, the stronger the vagal tone influence. For each day of evaluation and participant, we normalized the results computing the relative changes by subtracting the values of the experimental condition with that of the rest. Then, because significant differences between the two evaluation days did not emerge (t-test p> .05), we averaged the relative changes of the two days of evaluation creating a single RMSSD index.

#### 2.4.3 Electrodermal Activity

As further index of ANS response, the Electrodermal activity (EDA) was measured with the Empatica E4 wristband by means of two silver (Ag) plated electrodes located under the snap-fastener of the band and placed in the ventral inner wrist, lined up between the middle and the ring hand fingers. The device measured the EDA with a sampling rate of 4 Hz, dispensing an alternating current of 8 Hz with a measurement range between .01 and 100 microSiemens. For each participant and epoch (rest and experimental), the data were exported in .CSV with the E4 manager software and pre-processed via the EDA Explorer web-based tool (Taylor et al., 2015). This web app was developed for portable physiological device, such as the Empatica E4, consisting of a machine learning algorithm that divided the EDA signal in 5 seconds epochs and automatically detected and labeled the artefactual epochs given participants’ temperature, motion-based activity — both parameters simultaneously measured by the wristband together with the EDA — and standard settings (Taylor et al., 2015). The artefactual epochs were corrected via a spline interpolation algorithm and further analyzed with the MATLAB toolbox Ledalab (Version 3.4.9 – Benedek & Kaernbach, 2010). For each condition, a Continuous Decomposition Analysis was applied. This method decomposes the skin conductance signal in its continuous tonic and phasic activity (Benedek & Kaernbach, 2010). For our aims, the tonic EDA level was considered and used as a measure of the sympathetic nervous system tone. Then, for each day of evaluation and each participant, we normalized the tonic skin conductance index by subtracting the value of the experimental condition with that of the rest. Then, since we did not find significant differences between the two evaluation days (t-test p> .05), we averaged the relative changes of the two days of evaluation obtaining a single index.

#### 2.4.4 Subjective perception of CL

Participants self-assessed the CL subjectively experienced during each evaluation session answering to the question *“How intense was the mental workload you experienced in the previous two hours on a scale from 1 (not intense at all) to 5 (very intense)?”*. The wording and the scale were custom-made and adapted from the last item of the Eysink’s et al. (2009) cognitive load questionnaire, measuring the overall cognitive load intensity experienced by participants. For each participant, the scores were manually recorded by the experimenter. No significant differences were found between the two evaluation days (t-test p> .05). Thus, we computed an average between the two days of evaluation to obtain a single index of subjective CL.

### 2.5 Data Analysis

Data were inspected for possible univariate outliers, defined as the data points above and below 3 SD. The outlier removal is a necessary operation because they could distort results by altering the mean and the standard deviation (Cousineau & Chartier, 2010), especially in ecological research, where possible spurious variables cannot be controlled and where participants moved freely in the environment increasing the probability of collecting artifacts in the physiological measures. Cousineau and Chartier (2010) recommend to delete outliers without excluding participants, but using imputation techniques to replace the missing data. Accordingly, we found a small number of univariate outliers — nine, corresponding to the 3% of all data points — that were replaced using the K-Nearest Neighbours approach (Troyanskaya et al., 2001). This machine learning algorithm predicts each missing value by computing a weighted average of the closest n data points — Euclidean distances — to the missing observations. The imputation was made using Python’s (Version 3.8.8) scikit-learn toolbox (Version 0.24.1) with default settings (n_neighbors= 5). We performed the statistical analysis with the JASP software (Version 0.14.1). For each variable, we tested whether the data points approached the normal distribution. Except for the Conscientiousness and Emotional Stability scales of the TIPI that were negatively skewed (-2.19 and -2.68 z scores above normality, respectively) and the subjective perception of CL positively skewed (2.15 z scores above normality), the whole variables approached a normal distribution, meeting one of the main assumptions that allow us to use parametric statistical tests.

We preliminary performed an independent sample t-test to control whether the index of CL (subjective and objective) and the personality factors were comparable between males and females. Then, a Pearson’s correlation was made to control whether the two objective indexes of ANS — RMSSD and tEDA — were associated.

### 2.6 Hypothesis testing

With reference to *Hypothesis 1*, we performed two independent Pearson’s correlation tests comparing, respectively, the subjective perception of CL with the parasympathetic nervous system (vagal tone indexed by RMSSD) and the sympathetic nervous system (indexed by the tEDA) parameters. A co-activation of the indices of the sympathetic and parasympathetic nervous system, although found in the literature, was beyond the scope of our research because it was modulated by demographic factors that cannot be controlled in our small reference sample (Weissman & Mendes, 2021). However, this aspect has been investigated with a correlation test in the preliminary analysis.

With reference to *Hypothesis 2.1*, other two partial Pearson’s correlation with the same parameters and the TIPI scales of Extraversion, Agreeableness, Conscientiousness, Emotional Stability and Openness to Experiences as covariates, were performed to test whether the personality factors could confound the correlation between the subjective perception of CL and ANS activity.

Regarding *Hypothesis 2.2*, three independent linear regressions were performed to test whether the personality factors could predict the objective, psychophysiological and the subjective, self-reported measures of CL. In each analysis related to *Hypothesis 2.2,* the linear regression assumptions were met; the predictors did not show collinearity (Tolerance > 0.5, VIF < 2, Condition Index < 30), the residuals were normally distributed with mean 0 and did not significantly correlate with each other (Durbin Watson Test *p*> .05). The personality factors were set as predictors, while the subjective perception of CL, the RMSSD and the tEDA were used as dependent variables, respectively. In each linear regression, we used a backward deletion method to select the significant predictors. The analysis opened with all predictors inside the model. At each step, one predictor at a time was excluded, the one with the highest p-value and always greater than .10. Deleted predictors were no longer inserted into the model again. The cycle of analysis stopped when there were no more predictors to discard, or when all remaining predictors had a *p*< .10.

## 3. Results

The descriptive statistics of the variables^3^ are shown in Table 2.

**Table 1:** the descriptive of the psychophysiological and the behavioural index.

| **Psychophysiological index** | **Mean±SD** |
| --- | --- |
| RMSSD | 2.565±11.96 |
| Tonic EDA | .149±.56 |
| **Behavioural index** |  |
| Extraversion | 4.787±1.45 |
| Agreeableness | 4.963±1.17 |
| Conscientiousness | 5.770±1.02 |
| Emotional Stability | 5.057±1.24 |
| Openness to Experiences | 4.630±1.24 |
| Subjective cognitive load | 3.085±.51 |

### 3.1 Preliminary analysis

Two separate independent sample t-tests were performed to control whether the variables used in the analysis differed for gender. We found no significant differences between males and females in RMSSD (*t*(28) = .894, *p*= .379), tEDA (*t*(28) = .187, *p*= .853), Extraversion (*t*(28) = -.443, *p*= .661), Agreeableness (*t*(28) = -.458, *p*= .650), Conscientiousness (*t*(28) = -.354, *p*= .726), Emotional Stability (*t*(28) = -1.447, *p*= .159), Openness to Experiences (*t*(28) = -.237, *p*= .814) and subjective CL (*t*(58) = .254, *p*= .801).

Finally, a Pearson’s correlation was made between RMSSD and tEDA to control for possible associations. We did not find a significant correlation between these two measures of the ANS (*r*= -.273, *p*= .144).

### 3.2 Hypothesis 1: correlation between subjective and objective measures of CL

Two separate Pearson’s Correlations were performed to test the association between the subjective perception of CL with the vagal tone (RMSSD) and the sympathetic nervous system (tEDA) index, respectively. When we compared the vagal tone index with the subjective perception of CL, no significant correlation was found for the RMSSD (*r*= -.226, *p*= .230). Similarly, we did not find a significant correlation between the CL and the tEDA (*r*= .160, *p*= .399).

### 3.3 Hypothesis 2.1: correlation between subjective and objective measures of CL controlling by personality factors

We performed two separate Pearson’s Partial Correlations to associate the subjective perception of CL with the vagal tone and the sympathetic nervous system activity using as covariates the TIPI scales of Extraversion, Agreeableness, Conscientiousness, Emotional Stability and Openness to Experiences. In the first analysis, the subjective perception of CL assumed a significant negative correlation with the RMSSD (*r*= -.539, *p*= .005). Contrariwise, the subjective perception of CL did not correlate with the tEDA (*r*= .182, *p*= .384).

### 3.4 Hypothesis 2.2: prediction of personality factors with respect to subjective and objective measures of CL

Three linear regression analysis with the backward deletion method were performed setting as predictors the TIPI scales of Extraversion, Agreeableness, Conscientiousness, Emotional Stability and Openness to Experiences and as dependent variables the subjective perception of CL, RMSSD and tEDA, respectively.

The Agreeableness, the Emotional Stability and the Openness to Experiences personality factors explained significant amount of the variance in the subjective perception of CL, F (3,26) = 6.13, *p*= .003, R^2^= .414, R^2^ = .347. The regression coefficient (B= -.192, 95% CI[-.329, -.056]) indicated that an increase in the Agreeableness corresponded, on average, to a .192 points decrease in the subjective perception of CL (*p*= .008 – Fig. 2A). Similarly, an increase in the Emotional Stability (B= -.156, 95% CI[-.311, -.001]) corresponded to a .156 points decrease in the subjective perception of CL (*p*= .048 – Fig. 2B). Contrariwise, an increase in the Openness to Experiences (B= .255, 95% CI[.100, .410]) corresponded to a .255 points increase of the subjective perception of CL (*p*= .002 – Fig. 2C).

**Figure 2:**
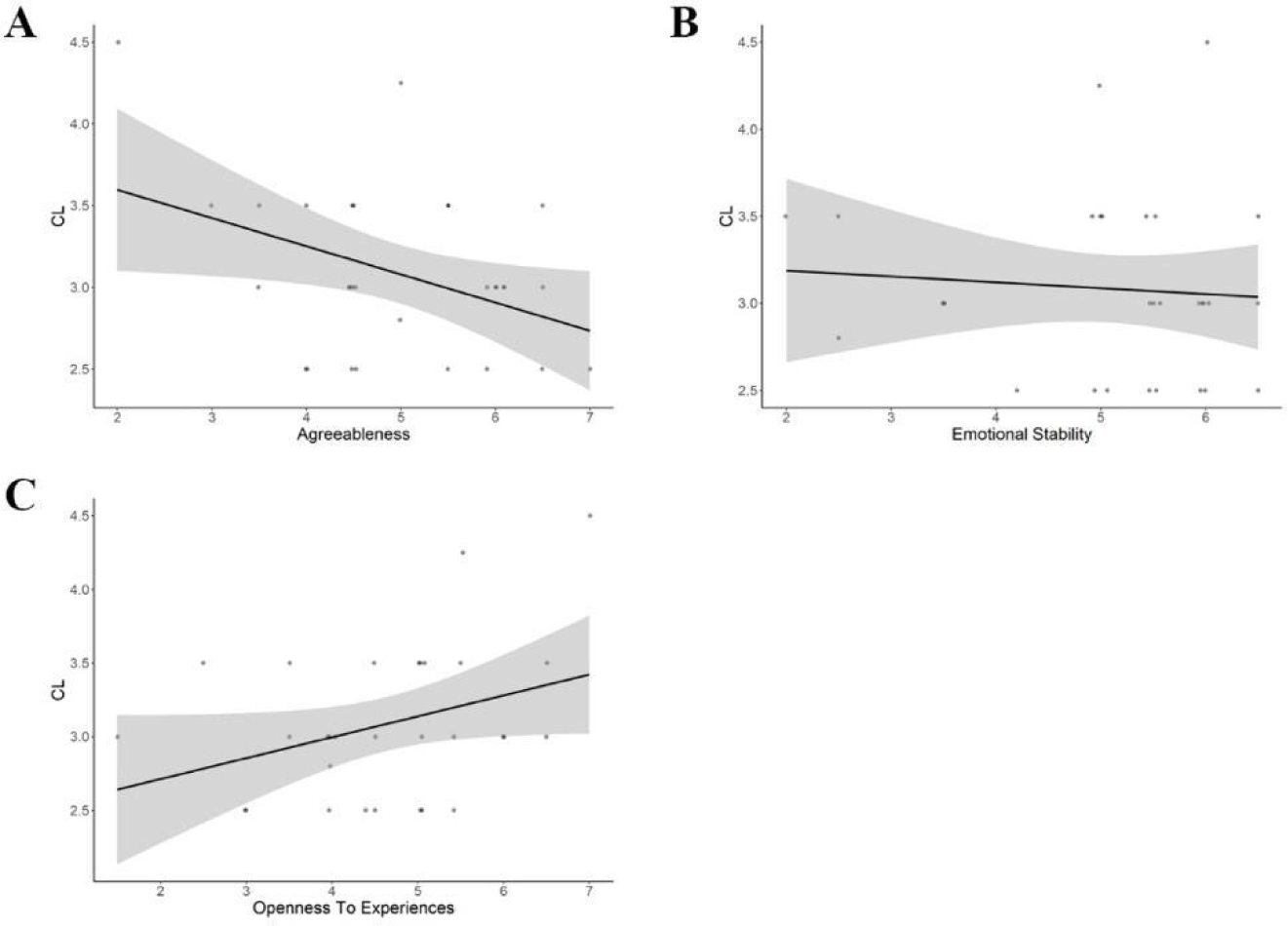
The scatterplot of the relation between the significant personality predictors of Agreeableness (A), Emotional Stability (B) and Openness to Experiences (C) and the dependent variable of subjective perception of the cognitive load (CL).

In the other linear regression analysis, the personality factors did not explain a significant amount of variance in the RMSSD, F (1,28) = 2.045, *p*= .164, R^2^= .068, R^2^ = .035, and in the tonic EDA, F (1,28) = 3.366, *p*= .077, R^2^= .107, R^2^ = .075.

## 4. Discussion

Our investigation explored a possible dissonance between the subjective and objective measure of CL, along with the influence of personality factors, on a population of workers (call center operators) in ecological conditions (standard working activities represented by inbound and outbound calls).

With reference to *Hypothesis 1*, we hypothesized to find no correlation between the subjective, self-reported measure of CL and its objective characterization, represented by the sympathetic and parasympathetic nervous system responses. In line with our expectations, no consistency was found between the subjective perception of CL and both the objective sympathetic and the vagal tone activity. Given the existing literature (e.g., Makransky, Terkildsen & Mayer, 2019; Minkley, Xu & Krell, 2021; Mutlu-Bayraktar, et al., 2020), the phylogenetic function of the sympathetic ANS is excitatory, mobilizing energies and preparing the organism to cope with a dangerous or stressful situation, such as increasing the heart and breathing rate. This system reflects a level of generalized activation of the organism (i.e., arousal) which increases in situations of (acute) stress. On the contrary, the parasympathetic ANS is an inhibitory system, which tends to restore the organism to a situation of calm and relaxation, for example by lowering the heart and breathing rate (Weissman & Mendes, 2021). Within this framework, our organizational population exhibits a lack of coherence between CL perception and both an overt stress expression and how stress is managed, and this lack of coherence represents possible evidence of CL dissonance. As discussed in the literature, call center operators are particularly prone to develop stress and other psychological problems (Conway et al., 2013; Costa & Costa, 2017; Jeyapal et al., 2015; Oh et al., 2017; Rameshbabu et al., 2013). In this vein, dissonance is instantiated in workers’ inability of openly express their emotional state of stress, which indeed is normally masked being substituted by a proper voice tone, and a calm attitude, during the interaction with customers (Molino et al., 2016; Zito et al., 2018). Our results show that this lack of coherence can emerge even during a typical working day. And, this could represent a risk for the workers’ well-being and health status, because it could favor the workers to underestimate the potential warning signals sent by their body or to overestimate their ability to manage stress. In fact, high CL levels in addition to other predisposing factors, including individual personality traits, are listed as some of the major sources of burnout (Koutsimani, Montgomery & Georganta, 2019; Maslach & Leiter, 2016). Different psychological training techniques were designed and experimented on call center operators to improve their “mind-body” coherence, such as mindfulness, defined as a practice that promotes a psychological state of awareness (Bossi et al., 2022; Davis & Hayes, 2011). Our evidence pushes even further towards the introduction of such techniques as a daily routine that organizational workers could practice during a typical work day to prevent possible health risks, such as CL dissonance (Bossi et al., 2022). Although outside of our aims, we cannot exclude that conditions of more extreme stress may highlight a correlation between subjective perception and ANS activation. Indeed, in our experimental task, we did not artificially induce high or low levels of stress or CL that could have taken to the extreme the scores of the variables and possibly revealed higher and significant associations. On the contrary, we studied participants in their standard and daily job activities, and in their ecological context. Moreover, we cannot rule out that a familiar setting, as compared to the laboratory, may have modified participants’ subjective and psychophysiological experience (Shamay-Tsoory & Medelsohn, 2019). In this vein, with reference to the embodied cognition perspective, there is increasing evidence that context can change our perception and the neural basis of our understanding (Willems & Peelen, 2021) and could be that in ecological, less extreme and less contrasting situations without considering possible confounding factors, the psychophysiological variables show some limits in predicting subjective experience of CL, and vice versa. Thus, future research should consider this methodological aspect also in investigating classical and well-established experimental topics.

With reference to *Hypothesis 2*, we explored how personality traits, measured through the TIPI, have played a confounding role in the association between the subjective and objective measures of CL, or predict their scores. Partially in line with our expectations (related to *Hypothesis 2.1*), we found that a correlation between the subjective perception of CL and the parasympathetic activity appears controlling for personality traits, while no correlation emerged with the sympathetic ANS. Specifically, the regression analysis (related to *Hypothesis 2.2*) revealed that the Openness to Experience, the Agreeableness and the Emotional Stability personality traits influenced the subjective score of CL. Contrariwise, no personality traits predicted the ANS measures. The personality trait of Openness to experience has been classically associated with a construct called need for cognition, i.e., the inclination to enjoy and engage in effortful though (Sadowski & Cogburn, 1997), as well of intellectual verbal skills and knowledge (Schretlen, van der Hulst, Pearlson & Gordon, 2010). Moreover, this trait has been positively associated with a high inclination to report high levels of stress in the workplace (Ervasti, Kallio, Määttänen, Mäntyjärvi & Jokela, 2019; Kim et al., 2016). Indeed, this trait was positively associated with CL perception in organizational workers. Agreeableness is a trait linked to prosociality, cooperation and generosity (Graziano & Eisenberg, 1997) and has been considered a protective factor for symptoms such as rumination, self-reported stress, depression and anxiety (Ervasti et al., 2019; Kim et al., 2016). Accordingly, in our participants, Agreeableness showed a negative association with subjective perception of CL. The Emotional Stability trait describes an emotionally stable, calm, relaxed and self-confident person, thus less prone to anxiety, moodiness, uneasiness or stress (Gosling et al., 2003). Accordingly, we found a negative association between CL perception and Emotional Stability.

Personality traits of Openness to Experience, Agreeableness and Emotional Stability have a direct relationship with CL perception and due to their direct influence on the subjective perception of CL, they could play a role as a confounding variable in the relationship between subjective and objective measures of CL (Figure 1B). Then, although we have no objective data about the actual workload that the participants carried out, we cannot exclude that these three personality traits may have boosted the coherence between the perceived CL and the intensity by which stress is managed, while they did not modify the lack of “mind-body” coherence with respect to the level of arousal. Commonly, in a typical working day, situations in which workers should manage many small stressful occurrences are more frequent compared to events of great intensity that cause high fluctuations in the body arousal. Thus, we can speculatively argue that, especially workers with a marked inclination both to report high levels of stress in the workplace (Openness to Experience) and less prone to experience anxiety, stress and depression (Agreeableness and Emotional Stability), are predisposed to manage and, accordingly, to focus their attention on how their own body is handling and coping to stressful situations. In other words, when perceived stress significantly increases or decreases, the workers might feel the need to manage their stress level to return to a state of equilibrium. Indeed, the vagal tone index, such as the RMSSD, is sensitive to small changes in cognitive activity related to mental fatigue and is associated with peoples’ ability to flexibly react to changing demands of the external environment (Matuz et al., 2021; Sebastiani et al., 2019). This skill is regulated by the inhibitory activity of the prefrontal cortex (Sebastiani et al., 2019) and in a typical working condition, the vagal tone may progressively reduce its activity in situations of acute stress, then regain vigor during a prolonged cognitive task which can cause mental fatigue (Matuz et al., 2021).

### 4.1 Limitations of the study

Our study presents five main limitations which should be considered for its generalization to other organizational settings.

First, our study was conducted on a restricted sample of workers (N=30). The limited sample size represents the trade off for an analysis conducted on real workers (call center operators) in strict ecological conditions (everyday, standard working activities, represented by inbound and outbound). Obviously, we fully acknowledge the importance and the reliability of larger experimental samples. However, we also want to mention that analyzing a population of real workers in natural conditions is not frequent in literature as the most of existing studies (albeit involving larger samples) are conducted on students facing laboratory tasks.

Second, we analyzed subjective and objective measures in standard, everyday working conditions, that is, we did not rely on an experimental design in which high *vs.* low CL were compared. The limited variability of the subjective and objective measures probably made our effect sizes less strong as compared to a laboratory context, in which more extreme stressful situations are deliberately induced. However, this limitation (as discussed previously) was also related to a deliberate theoretical choice. We aimed to investigate the actual activities of workers moving from the consideration that the sum of small daily stressful events can give rise to much more serious psychological problems over time. In other words, extreme stressful conditions, induced by the researcher, were not part of our analysis, considering also that extreme conditions could evoke incommensurable, hormesis-like, types of response (Calabrese & Baldwin, 2003).

Third, in our study we focused on CL, but the literature (discussed above) often refers to workload. Generally speaking, workload is not just related to trivial ‘levels’ of duties and ‘things to be done’, but to multiple characteristics of the task, which require cognitive integration (Chiorri et al., 2015b; Hancock et al.,1995; Hockey, 1997). In the existing literature, the preference for subjective measures of workload is a way to substitute the difficulty of objective measures. Among the subjective measures (e.g. Subjective Workload Assessment Technique, see Reid & Nygren, 1988), we opted for a simple item (5-points Likert scale) because of the presence of the limited amount of time and stringent ecological conditions. However, we are aware that measuring subjective CL related to workload is a still open issue, not part of our analysis.

Fourth, we acknowledge that the psychophysiological index that we used, the HRV and tEDA, are indirectly related to mental activity and can be influenced by various factors, not always related to the CL. Indeed, a brain activity index, like the EEG, could have given a more direct measure of CL. However, aimed to study workers in an ecological context over a few hours, we deliberately decided to use minimally invasive tools, such as a wearable wristband, in order not to significantly modify participants’ routine perception or create possible situations of discomfort. Equally, we adopted all the methodological procedures necessary to replace outliers, clean the physiological signal from artifacts or control the influence of other possible interfering variables, such as the increase in body temperature or body movements.

Fifth, in literature different validation studies were performed to investigate both the HRV and EDA reliability measure of the wristband we used. Indeed, the Empatica E4 is a device usable for research and non-clinical purposes, that was employed both in labs and natural environments, showing good reliability during rest, while low accuracy in dynamic conditions due to motion artifacts (Borrego, Latorre, Alcaniz & Llorens, 2019; Can, Arnrich & Ersoy, 2019; Milstein & Gordon, 2020; Stuyck, Dalla Costa, Cleeremans & Van den Bussche, 2022). Accordingly, we prevented the possibility of having a low signal-to-noise ratio acting both in the experimental design and in the data analysis phase. In fact, we mounted the device on the wrist of the left arm, which is less involved in participants’ work-related body movements. In addition, call center operators normally conduct their work activities seated in front of the desk, limiting possible uncontrolled large movements. Moreover, a thorough outlier detection and a re-entry of the missing data was carried out, also with the help of machine learning techniques that used acceleration and body temperature measures, collected in parallel by the device, for artifact detection.

## 5. Conclusions

When studying the complex relationship between subjective perception and objective experience of CL in organizational settings, it might be appropriate to consider workers’ psychological variables, such as personality traits, considered relatively stable and hardly modifiable factors of the human mind (Atherton, Grijalva, Roberts & Robins, 2020). Understanding the relationship between subjective and objective measures, and how it is confounded by personological traits present relevant *normative implications*, which could be of interest for both practitioners and workers.

With reference to practitioners (specifically managers and HR specialists), it is quite difficult to know in advance how much an employee is sensitive to changes in CL. Practitioners’ understanding is usually limited to the personological traits of the employees. Hence, knowing which personality traits may predict the individual perception of CL and which (consonant or dissonant) psychophysiological response is associated, could be very useful in practical terms. Speculatively such knowledge can be of help in designing ergonomics workplaces and promoting well-being practices in organizations.

With reference to workers, the eventual dissonance between subjective reports and objective physiological response could be a warning factor able to reveal a health risk, characterized by a discrepancy between worker’s troubled feelings and their “face” calm attitude. In particular, the inner experience of a stressful situation (instantiated in a specific physiological response) could be problematic when there is no correspondence with a subjective and conscious perception. Several training techniques are available, able to mitigate such kinds of uncomfortable situations in the workplace. Specifically, mindfulness represents techinque able to promote the awareness of mind-body integration (Davis & Hayes, 2011).

In conclusion, our study must be conceptualized within the contemporary trend of Organizational Neuroscience research, which is gradually moving from traditional ‘limited-to-the-brain’ analyses — conducted through the traditional techniques of brain imaging such as fMRI and EEG (e.g., Waldman, Balthazard & Peterson, 2011; for an overview see Becker, Cropanzano & Sanfey, 2011; Senior, Lee & Butler 2011) — to such investigations which take into account the fuller activities of the human nervous system involving the ANS (Massaro & Pecchia, 2019)^4^. An analysis of the ANS response (indexed by heart rate, pupil size and skin conductance) is able to shed light on the reliability and accuracy of more traditional constructs, based on subjective, self-reported perceptions, and to disentangle the role of personality traits in influencing CL.

## Footnotes

1 The ANS is part of the peripheral nervous system (composed by the cranial and spinal nerves), acts unconsciously and regulates involuntary activities such as digestion process, heart rate, pupil size, sexual arousal, etc. (for an overview, see Gabella, 2012). ANS is composed of the sympathetic and the parasympathetic nervous systems (notice that several authors consider the enteric nervous system as a third component). The sympathetic nervous system is associated with the ‘fight or flight’-type quick response and helps mobilize the body in demanding situations. The parasympathetic nervous system counteracts the sympathetic systems as it reinstates the homeostatic balance as being associated with ‘rest and digest’-or ‘feed and breed’-type responses (for an overview see Jänig, 2008).

2 The data that support the findings of this study are available on request from the corresponding author. The data are not publicly available due to privacy or ethical restrictions.

3 As discussed by Massaro & Pecchia (2019, p. 355) “measurements associated with the activity of the ANS, like cardiovascular measures, electrodermal activity (EDA or galvanic skin response), and blood pressure variation, have been largely disregarded” in organizational settings. Despite the existence of dedicated literature on the ANS in other branches, such as social psychology (e.g., Danyluck & Page-Gould, 2019), novel paradigms based on physiological variables can be of interest to organizational scholars.

4 A “confounder” or a “confounding variable” is not an intermediate factor between variable A and B, but an extraneous factor X that is independently associated both to variables A and B, thus potentially influencing the direction and size of the relation between A and B (Skelly, Dettori & Brodt, 2012). In our case, we hypothesize a confounding role of personality, because we found in literature that it is independently associated with both subjective and objective measures of cognitive load.

